# Anatomical Connectivity of the Macaque Gaze-following Patch

**DOI:** 10.64898/2026.10.27.762319

**Authors:** Ian Chong

## Abstract

The macaque gaze-following patch (GFP) is a specialized region of the temporal cortex implicated in decoding others’ focus of attention and guiding gaze-following behavior. Although the GFP has been characterized functionally, its anatomical connectivity remains largely unknown. To investigate how the GFP is embedded within a putative brain network supporting social interactions, we injected a pair of fluorescence labeled anterograde and retrograde tracers into the GFP of one monkey. Analysis of the resulting material revealed reciprocal connections between the GFP and a region of the inferior parietal lobule, likely corresponding to area PFG, as well as sparse retrograde labeling in the amygdala. In contrast to the expectation that the GFP might be subject to cognitive control signals from the prefrontal cortex, we failed to identify convincing retrograde labeling in prefrontal regions. Overall, this connectivity profile is consistent with a network that translates information derived from others’ faces into shifts of attention, while providing little evidence for direct prefrontal control of the GFP.

## 1. Introduction

When a human or a monkey tries to identify where another individual is looking at, a specific part of the posterior superior temporal cortex (pSTS) becomes activated (1,2). Single unit recordings from this cortical area in monkeys, called the gaze-following patch (GFP) have revealed neurons uniquely linking another’s gaze direction to specific spatial targets, and enabling flexible, context-dependent shifts of attention (3). Subsequent causal manipulation of this patch substantiated its role in gating gaze-following behavior through context, and led to the hypothesis that the GFP may form the core of a prefrontal-temporal-parietal network that governs when gaze-following should take place (4). The prefrontal cortex was suggested to be a viable source of top-down control signals that would instruct the GFP on when gaze-following behavior is appropriate or not, while the parietal cortex would be continuously updated by the GFP on where attention should be allocated and mediate the subsequent redirection of attention. More specifically the dorsal lateral prefrontal cortex (dlPFC) and the lateral intraparietal cortex (LIP) were proposed as respective up- and downstream connections with the GFP, as both brain areas seemed to fulfill promising anatomical (5,6) and physiological criteria (7,8).

Besides these two areas of interest, there are a multitude of other brain areas also worth considering for connectivity with the GFP. These include the face patch network, early visual cortical areas, the middle temporal visual area (MT), posterior dorsal inferotemporal cortex (PITd)/middle STS (mSTS), and subcortical areas such as the amygdala and superior colliculus, all of which may contribute or receive pertinent sensory, social and attention-related information and may be part of a larger joint attention network with the GFP (9–11).

Testing whether or not the above proposal is feasible requires the use of anatomical tracers to see which brain areas the GFP is connected to. This is not so easily accomplished considering the scarcity of monkeys with their GFP localized based on reliable electrophysiological criteria. The paradigms used to elicit gaze-following behavior for locating the GFP are not straightforward to train in monkeys, and as a result GFP monkeys with their namesake brain area fully characterized are rare. This is in stark contrast with the ease with which face patches can be identified in primates, as passive face perception tasks only require the maintenance of fixation as images are presented (12,13). We were fortunate to have in our possession a relatively aged rhesus macaque who had previously gone through both the GFP localization experiments via fMRI (1) as well as the single-cell electrophysiological experiments to characterize the GFP at the neuronal level (3). The GFP of this monkey had even undergone causal manipulation experiments to concretely tie this cortical patch to gaze-following behavior (4). This individual presented a unique opportunity to try to shed first light on the afferent and efferent connections of the GFP using retrograde and anterograde neuronal tracers.

The aims of the anatomical experiment in which we injected both an antero- and a retrograde tracer into the GFP were as follows: 1) clarify which part of the pSTS the macaque GFP is located in according to published macaque brain atlases, 2) look for labeling in the face patch system and nearby attention areas such as PITd, 3) check if there is anterograde and retrograde labeling of LIP and dlPFC, and 4) assess other brain areas that might be connected with the GFP that we did not include in a previously published connectivity model suggested by causal manipulation of the monkey GFP (4).

## 2. Methods

### 2.1 Animals and surgery

One male rhesus macaque (*Macaca* mulatta, age 18, 11kg) that had previously participated in functional MRI (1), electrophysiological (3) and reversible inactivation (4) studies to localize, characterize and demonstrate behavioral causality of the GFP was used for the present study (monkey L). Monkey L had been implanted with a titanium head-post for painlessly restraining the monkey’s head during experiments and a titanium chamber for accessing the STS. All experimental preparations and procedures were approved by the local animal care committee (Regierungspräsidium Tübingen, Abteilung Tierschutz) and were fully in line with German and European law and the National Institutes of Health’s Guide for the Care and Use of Laboratory Animals.

### 2.2 Tracer Injections

Tracer injections comprised of pressure injections made with glass micropipettes connected to a Hamilton syringe. Micropipettes were prepared from glass capillaries (outer diameter 0.5mm, inner diameter 0.33mm, Hilgenberg) via a P-30 Micropipette puller (Sutter Instruments, Novato). The glass micropipette was glued to a custom adaptor allowing airtight connection to a modified 10μl Hamilton syringe. Both were prefilled with paraffin oil before being inserted into a 20G metal cannula. 1μl of the tracer of interest was loaded into the micropipette from the tip under a dissecting microscope, and the micropipette was withdrawn 1cm away from the sharp opening of the cannula before penetrating the dura.

The injection site of the tracer corresponded to the ‘GFP hotspot’ characterized by the highest contrast between gaze-following and identity-mapping neurons (3), and the strongest gaze-following impairments following microstimulation and muscimol injections respectively (4). The entire micropipette/Hamilton syringe assembly was driven into the STS at 12μm/s via a micropositioner (David Kopf Instruments Model 650) until it reached the target depth (approximately 5mm under the dura to target the GFP, based on estimates from our prior recordings and reversible inactivation). The micropipette was allowed to stabilize for 15mins after the micropipette had reached the target depth before we commenced injections. 0.2μl of each tracer were injected across 45 minutes, with 30 minutes of waiting time after injection to allow full tracer uptake into the brain tissue before the pipette was slowly retracted out of the cortex.

The retrograde tracer cholera toxin subunit B (CTB)-Alexa Fluor 594 (1% PBS, C-34777, Invitrogen) and the anterograde tracer dextran-Alexa Fluor488 (10% PBS, D-22910, Invitrogen) were injected on different days into the right GFP of monkey L. Injections of CTB and dextran were made 14 and 15 days prior to the day of perfusion respectively.

### 2.3 Perfusion and Tissue Processing

On the day of the perfusion, anesthesia was initiated with ketamine (10mg/kg), followed by an overdose of barbiturate. Transcardial perfusion was performed under compete analgesia and unconsciousness. The perfusion was performed first with 3L of chilled (4°C) 0.12M phosphate buffer (PB) mixed with the anticoagulant Fraxiparine (5700 UI/L) to wash out blood, followed by 2L of chilled 0.12M PB mixed with 4% paraformaldehyde (PFA), and finally with 1L of chilled 0.12M PB mixed with 4% PFA and 10% sucrose. After the perfusion was completed, the brain was extracted from the skull and stored at 4°C in 1L of 4% PFA and 10% sucrose in 0.12M PB to begin the cryoprotection process. As soon as the brain had sunk to the bottom of the container after 1 day, we replaced the brain immersion bath with 1L of 4% PFA and 30% sucrose in 0.12M PB, and waited again for the brain to sink to the bottom of the container.

Once cryoprotection was complete, we immediately proceeded to cutting with a large-object cryotome (Leica) with cooled specimen stage and knife. The brain was coated in Tissue-Tek (Sakura) and dry ice fragments helped to speed up the cooling process. Once freezing was complete, sagittal 60μm sections were cut. We produced six series, of which only 2 were used for imaging. The brain sections were rinsed in floating solution (for 1L, 1g gelatin in 500ml distilled water, 400ml 95% ethanol, 75ml PB), mounted on gelatin-coated glass slides, air dried, cover slipped with mounting medium (DNA-SCR-038448, Biozol), and stored at 4°C in the dark.

### 2.4 Imaging and Data Analysis

Low power photomicrographs of entire sagittal slices were first generated using a Leica DMi8 fluorescence microscope (Camera: Leica DFC9000 GTC sCMOS, light source: Lumencor Spectra X 470 nm and 550/575 nm, filters: Alexa Fluor 488 and 555) and Las X software. Once retrograde and anterograde labeling were identified, higher power photomicrographs of these areas of interest were produced. Images were first processed on Las X for brightness and contrast, before being exported and processed in Adobe Photoshop 7.0.

The relatively large injection halo probably exceeded the boundaries of the GFP. Hence, the possibility of tracer uptake by neighboring areas, in particular the closely adjoining middle and posterior temporal face patches has complicated the interpretation of the labeling result in these areas. Unfortunately the interpretation of our tracing results was also severely impeded by the presence of strong auto-fluorescence from lipofuscin. Lipofuscin is known to accumulate over life, primarily in particularly metabolically active neurons (14,15) and was therefore unsurprisingly abundant in monkey L, who was 18 years old by the date of termination. Lipofuscin has an emission spectrum that is relatively broad and overlapping with the filters we used. Hence, we adopted strict selection criteria in the interpretation of our results to avoid reporting false labeling of neurons. Firstly, for retrograde labeling labeled cell bodies harboring the red fluorescent signal were required to be confined to specific cortical layers and present across adjacent brain slices. However this criterion was not sufficient as layer 5 houses the largest and metabolically most active pyramidal cells and is therefore arguably particularly prone to lipofuscin accumulation. This is why we resorted to a second criterion that required true labeling to be localized and moreover, be confined to subsets of sparse cells of similar size in a particular layer and region. Finally, labeled cells had to be sufficiently far apart from the original injection site in order to rule out neurons that had taken up tracer spilling over from the injection site.

## 3. Results

Here, we share preliminary tracing results that reveal for the first time the connectivity of the gaze-following patch (GFP) with the rest of the macaque brain. Although our results are based on data from one monkey only and therefore need to be substantiated in others, given the uniqueness of our sample we feel compelled to share our findings hoping that they may guide future work. Observations not fulfilling the strict criteria described in Methods to avoid false labeling due to lipofuscin are not reported.

Both the anterograde and retrograde injection site haloes place the GFP in the part of TEO that is located at the transition between the fundus and the lower bank of the posterior STS (corresponding to Figure 148 in (16)). This is in accordance with the location of the GFP in the original fMRI study (1) to which this monkey contributed. This position is anatomically distinct from areas MT/MST, and FST, which lie deeper in the fundus toward the lunate sulcus, and from PITd/IPa which is located more anteriorly in the middle STS (17). This allows us to conclude that the GFP is not part of any of these aforementioned brain areas, and moreover based on location alone is unlikely to be one of the classical face patches.

### 3.1 Anterograde projections of the GFP to other parts of the STS and early visual cortex

Anterograde labeling originating from the GFP extended several millimeters beyond the injection halo into neighboring regions of the posterior STS. Labeled axons and terminal-like structures were observed in adjacent cortical areas, including regions corresponding to early visual areas V2, V3, and V4, as well as PITd. Labeling was also present along the lower bank and fundus of the posterior STS, extending both anteriorly and posteriorly from the injection site.

Given the proximity of the middle and posterior face patches to the GFP, and the sensitivity of the middle face patches (ML and MF) to head direction (11,18,19), the GFP is well positioned to receive face—and possibly eye—direction information from the face-patch system, as proposed in our gaze-following model (4). Monkey L was part of the fMRI experiments that first localized the GFP and canonical face patches (1), allowing us to estimate face-patch locations in the histological sections. Although anterograde labeling was observed in these regions, the labeled axons appeared continuous with those surrounding the injection site rather than forming discrete terminal fields, and no clear labeling was detected in the estimated MD face area. Thus, the broad, continuous labeling likely reflects projections to neighboring posterior STS regions. Because the tracer may have extended beyond the GFP boundaries, however, we cannot confidently attribute labeling in adjacent face-patch areas to GFP projections.

### 3.2 Reciprocal connectivity between the GFP and the posterior parietal cortex

Contrary to the expectations prompted by our previous physiological work, we were unable to find retrograde or anterograde labeling in area LIP and dlPFC. Instead, we discovered both retrograde and anterograde labeling in a restricted part of the posterior parietal cortex (Fig. 2), most likely corresponding to area PFG, consistently observed across multiple adjacent sagittal sections. Retrograde labeled cell bodies were confined to layer V (Fig. 2B), whereas anterograde terminal axonal labeling was concentrated within the superficial cortical layers (I-III) (Fig. 2A). In contrast to several other cortical regions displaying widespread lipofuscin-based auto-fluorescence, the labeling in this region was spatially restricted, exhibited clear laminar organization, and fulfilled our predefined criteria for accepting tracer labeling. We therefore interpret these observations as evidence for reciprocal anatomical connections between the GFP and the posterior parietal cortex.

**Figure 1.**
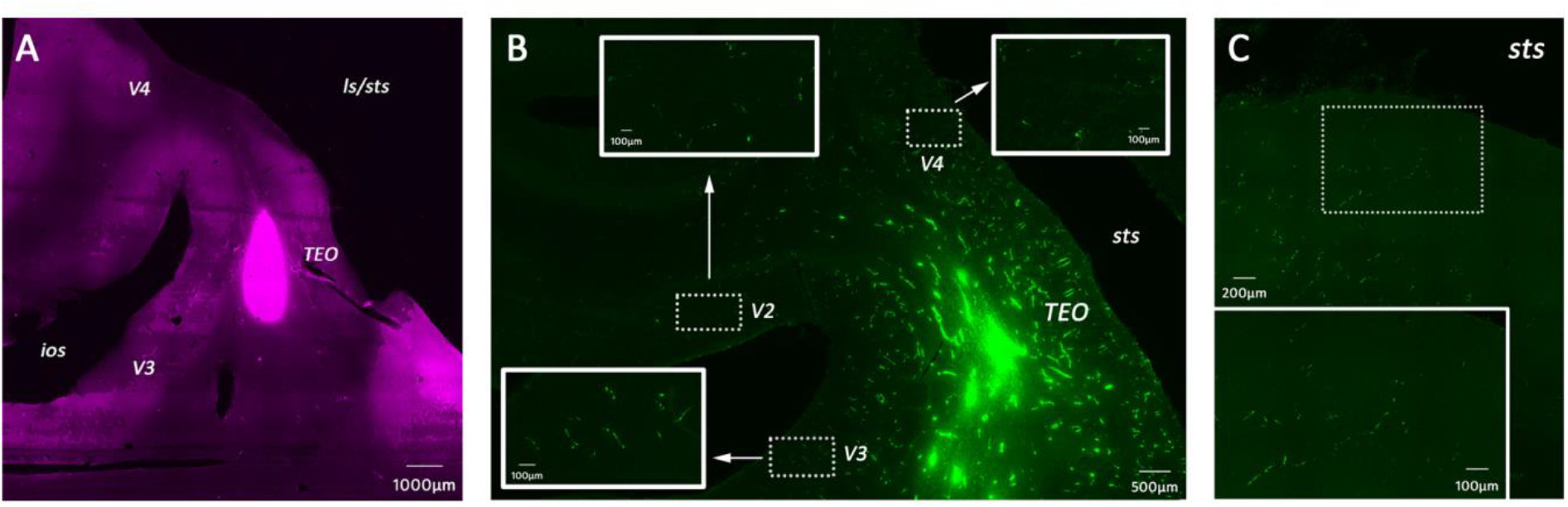
Tracer injection halo and local anterograde connectivity. A. Fluorescent micrograph of the injection halo of the retrograde tracer CT-B-Alexa Fluor594 on a sagittal slice, corresponding to the GFP. B. Local connectivity of the anterograde tracer dextran-Alexa Fluor488. The injection halo of the tracer is clearly visible and overlaps with the halo shown in A. Insets (white boxes) correspond to sections of the micrograph (dotted boxes) indicated by the white arrows. Local anterograde labeling was found in the vicinity of the injection halo (4-5mm away), including early visual areas V2, V3, and V4. C. Example of unspecific anterograde labeling found in expected location of the face patches based on the subject’s (M1) fMRI data (1); in this case the location of face patch MF is shown. Abbreviations: ios, inferior occipital sulcus; ls, lateral sulcus; TEO, temporo-occipital cortex; sts, superior temporal sulcus.

**Figure 2.**
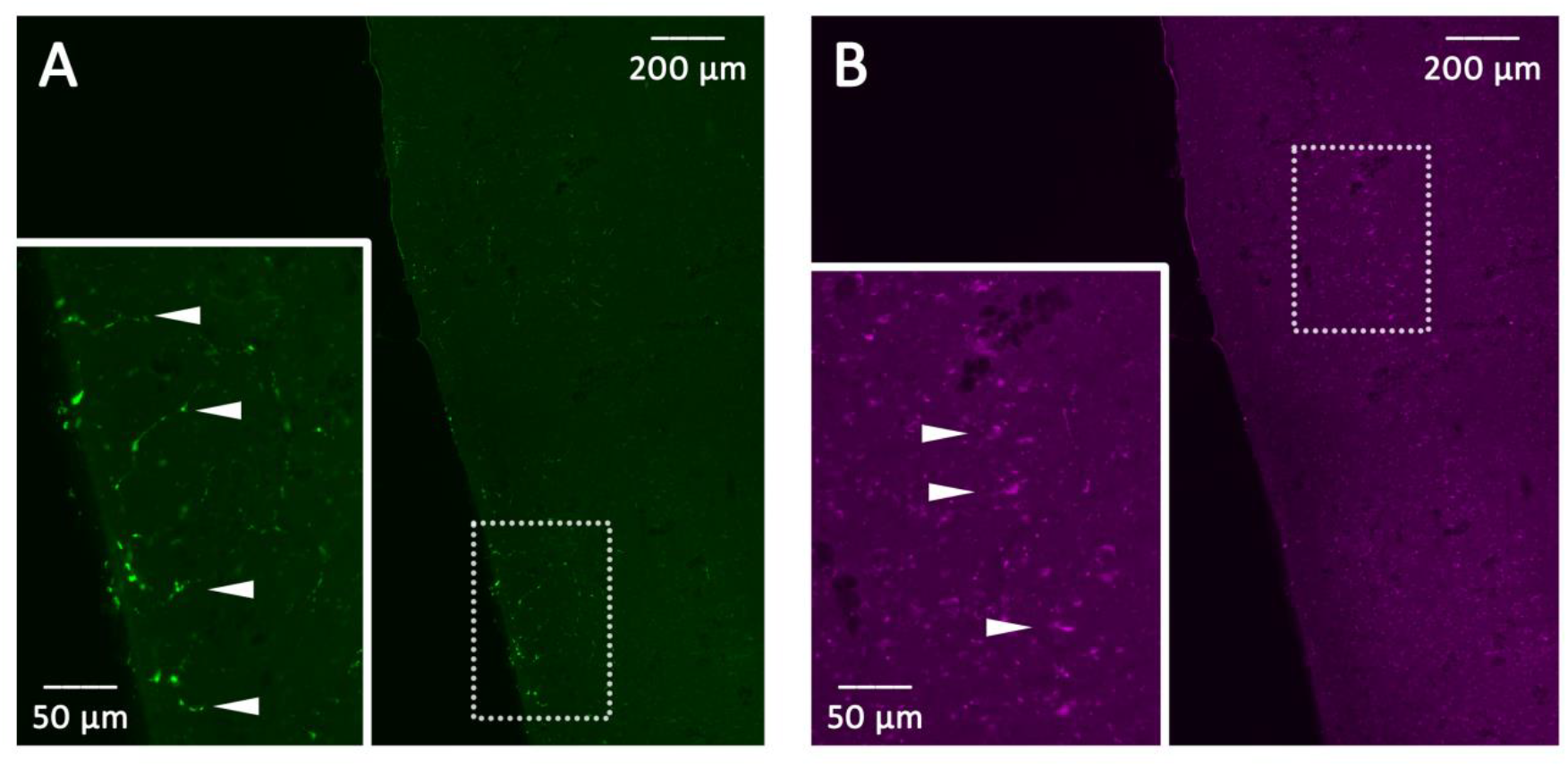
Anterograde and retrograde labeled cells in the posterior parietal cortex. A. Anterograde labeling was found in area PFG. Labeled cells run along the entire edge of the displayed section, and the inset shows a magnified view of the area within the dotted box. White arrows point out specific terminal axon labeling that was restricted to superficial cortical layers (I-III). B. Retrograde labeling was found in the same area (PFG) as the anterograde labeling but in layer V. White arrows point out labeled cell bodies, which trail along the entire length of the displayed section. The inset shows a magnified view of the area within the dotted box.

### 3.3 Afferent connectivity from the amygdala to the GFP

Among the subcortical structures examined, only the amygdala exhibited convincing retrograde labeling. Clear labeling of individual, sparsely distributed cell bodies could be identified (Fig.3). The labeling we saw could be found in adjacent sagittal brain slices and were mostly found in the basolateral amygdala.

### 3.4 Unresolved retrograde labeling of the motor, premotor, and somatosensory areas

Unexpectedly we discovered what appeared to be retrograde labeling in the primary motor cortex F1, premotor cortex F2, primary somatosensory cortex, and parts of the anterior and posterior cingulate cortex. Although the fluorescent retrograde signal was intense and confined to layer V, the labeling failed to meet the criterion that only sparse subsets of cells in restricted regions should exhibit tracer signals. For example, in the primary motor cortex, labeled cell bodies were distributed broadly from the lateral to the medial side in layer 5 (20).

To fully convince ourselves that these observations reflected age-related auto-fluorescence rather than true tracer transport, we examined brain tissue from an age-matched macaque that had not been subject to tracing experiments, and looked for stereotypical patterns of auto-fluorescence. Indeed a similar pattern of auto-fluorescence was present in the same cortical regions highlighted above, indicating that these signals were most likely attributable to lipofuscin accumulation rather than retrograde labeling. Additional areas that also harbored tremendous amounts of auto-fluorescence in both the experimental animal and the control included the lateral geniculate nucleus (LGN) and the globus pallidus (GP). Consequently, labeling in these cortical and subcortical regions was considered inconclusive and excluded from further interpretation.

## 4. Discussion

The present study represents the first anatomical investigation of the cortical and subcortical connectivity of the macaque gaze-following patch (GFP). Unfortunately, extensive lipofuscin auto-fluorescence associated with the age of the monkey used in this study has confounded the interpretation of our results, and therefore only labeling that fulfilled extremely stringent anatomical criteria was considered as trustworthy. Despite this technical limitation, we identified convincing reciprocal connectivity with a restricted region of the posterior parietal cortex, corresponding to area PFG (Fig. 2). Moreover, we also observed subcortical input originating from the basolateral amygdala (Fig. 3). By contrast, we found no convincing evidence for direct connectivity with several nodes in our previously published network model of gaze-following suggested by functional data. In particular, we failed to identify connections with dorsolateral prefrontal cortex and area LIP. On the other hand, we obtained evidence for input to the GFP from the basolateral amygdala, previously not foreseen in the aforementioned model. Yet, considering its well-recognized role in evaluating socially relevant information, including facial expressions, eye contact, and gaze direction this connection might have been expected (21,22). Recalling that gaze-following can be influenced by facial expressions (23–25) and direct gaze (23), arguably processed in the amygdala, suggests a potential pathway through which the modulation of gaze-following by social context could be achieved.

**Figure 3.**
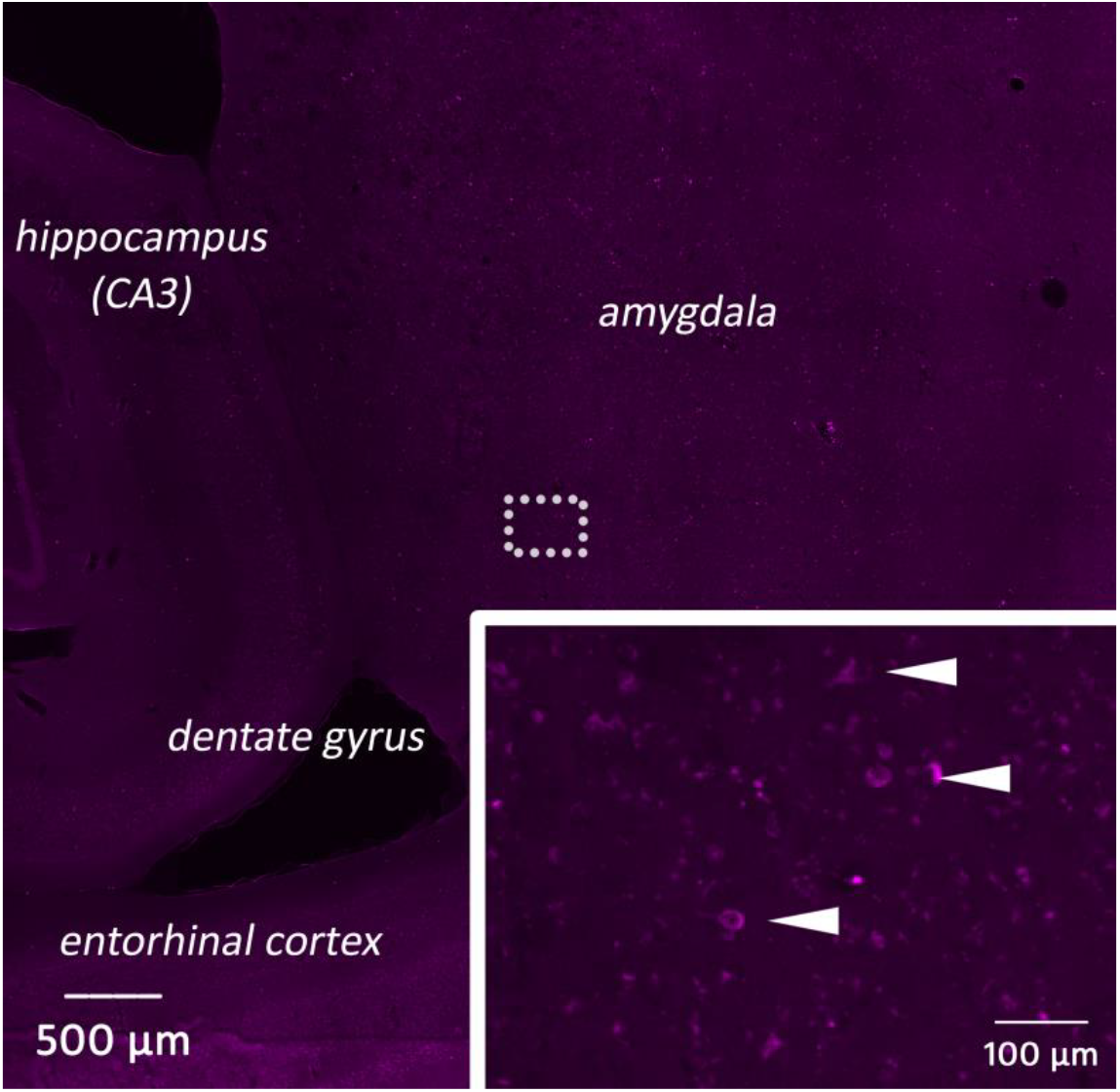
Subcortical retrograde labeling in medial temporal lobe structures. Sparse retrograde labeling was identified in the amygdala. Inset shows highlighted region in the dotted area, and arrows indicate labeled cell bodies.

The most robust long distance tracing result of the present study was the reciprocal connection between the GFP and area PFG (Fig. 2). This finding was unexpected in light of our previous block model, which posited interactions between the GFP, LIP, and prefrontal cortex during gaze-following (4). Instead, our preliminary data points to the inferior parietal lobule as a prominent cortical partner of the GFP. Connections between the STS and PFG have been reported previously in macaque tracing studies (26,27). Additional tracing studies have described projections from the posterior parietal cortex into the lower bank of the STS (28,29), which may potentially overlap with the GFP. Thus, although the GFP itself has not been previously examined anatomically, the present findings are broadly consistent with known STS-parietal connectivity patterns.

The finding that no connections could be found between the GFP and LIP is unexpected because the latter area has been implicated in the selection of spatial targets singled out by gaze cues (8) and in the control of visually guided saccades; functions that appear particularly relevant for gaze-following in which another’s gaze is translated into a spatially directed eye movement. Area PFG contains neurons that respond during both the execution of an action and the observation of others performing the same action, i.e. mirror neurons. The current anatomical finding raises the possibility that another individual’s gaze may be integrated with representations of their actions or behavioral goals. During natural social interactions, gaze cues rarely occur in isolation, and where another is looking can be informative of goal-directed behavior towards a behaviorally relevant object (30). Reciprocal communication between the GFP and PFG could therefore provide a pathway through which gaze information contributes to the understanding of observed behavior (27) and the selection of an appropriate response. This interpretation also provides a possible functional context for the role of gaze-following itself. Following another individual’s gaze allows an observer to orient towards objects that are relevant to the other, but the resulting information can subsequently be used to guide one’s own behavior too. A connection between the PFG and the GFP could therefore link gaze-following with parietal mechanisms involved in action observation and goal-directed behavior. In this framework, the GFP would not necessarily function as part of a gaze-to-saccade pathway, but would be part of a social perception network where gaze direction can be used to infer behavioral significance of the other’s actions. This framework is backed by studies showing that the STS and inferior parietal lobule where PFG is located are involved in the observation of actions (27,31), actions with intention (32) and more broadly social interactions (33).

Finally, we were unable to identify any connections with prefrontal areas, which were previously hypothesized to be a potential source of top-down cognitive control signals impacting the GFP. This does not necessarily falsify the hypothesis that top-down cognitive control signals instructing gaze-following originate from the prefrontal cortex. A possible explanation might be insufficient transport of tracer due to too short survival times, or a polysynaptic pathway rather than a direct connection. In fact, this result is similar to the report that the anterior face patches in the inferotemporal (IT) cortex receive only weak inputs from prefrontal cortex (34), in contrast to the expectation that area TE –housing these patches– should receive many projections from prefrontal areas (35).

We end this study with a final word of caution, emphasizing the preliminary nature of the findings and their interpretations. First of all, one should keep in mind that the results reported are from one individual only and are in urgent need of confirmation in more animals. While the injections carried out were guided by physiological data on the location of the GFP, only the congruency of injection patterns in more animals will alleviate concerns of unintended spread of the tracer from the GFP injection site. New protocols also need to be designed to locate the GFP in non-human primates more swiftly without resorting to our previously used gaze-following paradigm that requires lengthy training (1,3,4). With improved methods to identify the GFP in the primate brain we will be able to circumvent the use of older animals who had to undergo years of testing, and avoid the glaring issue of lipofuscin buildup. Secondly if future experiments would resort to the same tracers, the volumes of injections should be further reduced and the survival times –decisive for the distance tracers can cover – be varied. Finally, given the high likelihood of substantial auto-fluorescence even in somewhat younger animals, which may compromise the reliable assessment of labeling, tracer detection should rely on immunohistochemistry rather than fluorescence tagging.

## Acknowledgements

I thank Professor Peter Thier for his supervision in this pilot project, and providing helpful feedback in preparing this report.

## References

1. Marciniak K, Atabaki A, Dicke PW, Thier P. Disparate substrates for head gaze following and face perception in the monkey superior temporal sulcus. Elife. 2014;3(July 2014):1–18.

2. Marquardt K, Ramezanpour H, Dicke PW, Thier P. Following eye gaze activates a patch in the posterior temporal cortex that is not part of the human “face patch” system. eNeuro. 2017;4(2):1–10.

3. Ramezanpour H, Thier P. Decoding of the other’s focus of attention by a temporal cortex module. Proc Natl Acad Sci U S A. 2020;117(5):2663–70.

4. Chong I, Ramezanpour H, Thier P. Causal manipulation of gaze-following in the macaque temporal cortex. Prog Neurobiol. 2023 Jul;226(December 2022):102466.

5. Baizer JS, Ungerleider LG, Desimone R. Organization of visual inputs to the inferior temporal and posterior parietal cortex in macaques. J Neurosci. 1991;11(1):168–90.

6. Yeterian EH, Pandya DN, Tomaiuolo F, Petrides M. The cortical connectivity of the prefrontal cortex in the monkey brain. Cortex. 2012;48(1):58–81.

7. Wallis JD, Anderson KC, Miller EK. Single neurons in prefrontal cortex encode abstract rules. Nature. 2001;

8. Shepherd S V., Klein JT, Deaner RO, Platt ML. Mirroring of attention by neurons in macaque parietal cortex. Proc Natl Acad Sci U S A. 2009;106(23):9489–94.

9. Leopold DA, Krauzlis RJ. How the brain pays attention to others’ attention. Proceedings of the National Academy of Sciences of the United States of America. 2020.

10. Bogadhi AR, Bollimunta A, Leopold DA, Krauzlis RJ. Spatial Attention Deficits Are Causally Linked to an Area in Macaque Temporal Cortex. Curr Biol. 2019;29(5):726–736.e4.

11. Yang Z, Freiwald WA. Joint encoding of facial identity, orientation, gaze, and expression in the middle dorsal face area. Proc Natl Acad Sci U S A. 2021;118(33).

12. Tsao DY. A Cortical Region Consisting Entirely of Face-Selective Cells. Science (80-). 2006 Feb;311(5761):670–4.

13. Hesse JK, Tsao DY. The macaque face patch system: a turtle’s underbelly for the brain. Nat Rev Neurosci. 2020;21(12):695–716.

14. Terman A, Brunk UT. Lipofuscin. Int J Biochem Cell Biol. 2004 Aug;36(8):1400–4.

15. Brunk UT, Terman A. Lipofuscin: mechanisms of age-related accumulation and influence on cell function. Free Radic Biol Med. 2002 Sep;33(5):611–9.

16. Saleem KS, Logothetis NK. Saleem KS, Logothetis NK. A combined MRI and histology atlas of the rhesus monkey brain in stereotaxic coordinates. Academic Press; 2012.

17. Stemmann H, Freiwald WA. Evidence for an attentional priority map in inferotemporal cortex. Proc Natl Acad Sci U S A. 2019;116(47):23797–805.

18. Taubert J, Japee S, Murphy AP, Tardiff CT, Koele EA, Kumar S, et al. Parallel processing of facial expression and head orientation in the macaque brain. J Neurosci. 2020;40(42):8119– 31.

19. Freiwald WA, Tsao DY. Functional Compartmentalization and Viewpoint Generalization Within the Macaque Face-Processing System. Science (80-). 2010 Nov;330(6005):845–51.

20. Graziano MSA. Ethological Action Maps: A Paradigm Shift for the Motor Cortex. Trends Cogn Sci. 2016;20(2):121–32.

21. Mosher CP, Zimmerman PE, Gothard KM. Neurons in the monkey amygdala detect eye contact during naturalistic social interactions. Curr Biol. 2014;24(20):2459–64.

22. Minxha J, Mosher C, Morrow JK, Mamelak AN, Adolphs R, Gothard KM, et al. Fixations Gate Species-Specific Responses to Free Viewing of Faces in the Human and Macaque Amygdala. Cell Rep. 2017;18(4):878–91.

23. Chong I, Thier P. Monkeys integrate facial expressions and direct gaze to modulate gaze-following behavior. Curr Res Neurobiol [Internet]. 2026 Jun;10(June 2025):100158. Available from: https://linkinghub.elsevier.com/retrieve/pii/S2665945X26000045

24. Teufel C, Gutmann A, Pirow R, Fischer J. Facial expressions modulate the ontogenetic trajectory of gaze-following among monkeys. Dev Sci. 2010;13(6):913–22.

25. Goossens BMA, Dekleva M, Reader SM, Sterck EHM, Bolhuis JJ. Gaze following in monkeys is modulated by observed facial expressions. Anim Behav. 2008;75(5):1673–81.

26. Rozzi S, Calzavara R, Belmalih A, Borra E, Gregoriou GG, Matelli M, et al. Cortical Connections of the Inferior Parietal Cortical Convexity of the Macaque Monkey. 2006;(October):1389–417.

27. Nelissen K, Borra E, Gerbella M, Rozzi S, Luppino G, Vanduffel W, et al. Action observation circuits in the macaque monkey cortex. J Neurosci. 2011;31(10):3743–56.

28. Neal JW, Pearson RCA, Powell TPS. The connections of area PG, 7a, with cortex in the parietal, occipital and temporal lobes of the monkey. Brain Res. 1990;532(1–2):249–64.

29. Neal JW, Pearson RCA, Powell TPS. The cortico-cortical connections within the parietotemporal lobe of area PG, 7a, in the monkey. Brain Res. 1988;438(1–2):343–50.

30. Shepherd S V. Following gaze: Gaze-following behavior as a window into social cognition. Front Integr Neurosci. 2010;4(MARCH 2010):1–13.

31. Molenberghs P, Brander C, Mattingley JB, Cunnington R. The role of the superior temporal sulcus and the mirror neuron system in imitation. Hum Brain Mapp. 2010;31(9):1316–26.

32. Di Cesare G, Lombardi G, Zeidman P, Urgen BA, Sciutti A, Friston KJ, et al. Two distinct networks for encoding goals and forms of action: An effective connectivity study. Proc Natl Acad Sci. 2024 Jun;121(26):1–7.

33. Sliwa J, Freiwald WA. Neuroscience: A dedicated network for social interaction processing in the primate brain. Science (80-). 2017;356(6339):745–9.

34. Grimaldi P, Saleem KS, Tsao D. Anatomical Connections of the Functionally Defined “Face Patches” in the Macaque Monkey. Neuron. 2016;90(6):1325–42.

35. Borra E, Ichinohe N, Sato T, Tanifuji M, Rockland KS. Cortical connections to area te in monkey: Hybrid modular and distributed organization. Cereb Cortex. 2010;20(2):257–70.

